# Aim11 is a novel protein involved in the assembly of mitochondrial cytochrome *c* oxidase

**DOI:** 10.64898/2025.12.16.694746

**Authors:** Ulrik Pedroza-Dávila, Yolanda Camacho-Villasana, Miriam Vázquez-Acevedo, Madhurya Lutikurti, Diego González-Halphen, Alfredo Cabrera-Orefice, Xochitl Perez-Martinez

## Abstract

Cytochrome *c* oxidase (CIV) is the last electron acceptor of the mitochondrial respiratory chain. In yeast, it is composed of 12 subunits, three of which are encoded in the mitochondrial genome. CIV assembly is a modular and highly regulated process that requires several specific factors. In this work, we characterized the role of Aim11 in CIV biogenesis. By high-throughput analysis, it was previously detected that Aim11 interacted with some CIV subunits, but the physiological relevance of these interactions was unknown. In the present work, we found that the *Δaim11* mutant exhibited reduced respiratory growth and diminished CIV activity. Using mitochondrial complexome profiling, we detected in the *Δaim11* mutant accumulation of intermediates of the three CIV-assembly modules, as well as reduction in supercomplexes levels. Aim11 works together with three other uncharacterized proteins: Mtc3, Gep7, and Iai11. The four proteins form a complex that we named AMIGa (Aim11-Mtc3-Iai11-Gep7 association) complex, necessary for the efficient assembly of CIV. Finally, the human protein TMEM242 was identified as Aim11 orthologue.

## INTRODUCTION

The oxidative phosphorylation system (OXPHOS) comprises the respiratory complexes and ATP synthase. Cytochrome *c* oxidase (Complex IV, CIV) is the last electron acceptor of the mitochondrial respiratory chain and is responsible for donating electrons to molecular oxygen. In yeast mitochondria, it is composed of three core subunits (Cox1-3) whose genes are in the organelle’s genome (mtDNA) and nine accessory subunits (Cox4-9, Cox12, Cox13, and Cox26) encoded in nuclear genes. Due to their dual genetic origin, the assembly of CIV subunits must be coordinated to ensure correct biogenesis and stoichiometry (Moretti-Horten *et al*, 2024). The assembly of CIV is proposed to be modular, where each mtDNA-encoded subunit is associated with a specific set of accessory subunits and assembly factors and then the three distinct modules assemble to produce the mature CIV(Su *et al*, 2014b; Franco *et al*, 2018; McStay *et al*, 2013b). However, it remains unknown how some of these biogenesis steps occur and which proteins are involved in the regulation of these processes.

In this work, we characterized the mitochondrial protein Aim11 and unveiled its role in the assembly of complex IV. Aim11 is a mitochondrial inner membrane protein of ∼15 kDa with two transmembrane helices (Morgenstern *et al*, 2017). Aim11 loss was shown to be lethal in the absence of prohibitins (Osman *et al*, 2009). Additionally, a large-scale pulldown experiment showed that Aim11 physically interacts with some CIV subunits and assembly factors, such as Cox2, Cox20, and Cox6 (Morgenstern et al., 2017). However, the relevance of these interactions was not established. Here, we found that Aim11 is necessary for the optimal assembly of complex IV, affecting its three assembly modules (Cox1-3). We show that Aim11 forms a complex with the uncharacterized proteins Iai11, Gep7, and Mtc3, and we named it AMIGa (Aim11-Mtc3-Iai11-Gep7 association) complex. All four components are necessary for optimal CIV assembly. Finally, we identified TMEM242 as the human functional orthologue of Aim11, which was able to compensate for the loss of Aim11 in yeast.

## RESULTS AND DISCUSSION

### Aim11 is involved in CIV biogenesis

By high-throughput assays, it was previously observed that Aim11 co-purified with some CIV subunits (Morgenstern *et al*, 2017); however, the physiological relevance of these interactions remained unexplained. To explore the role of Aim11 in CIV biogenesis, we characterized a *Δaim11* mutant. First, we performed serial dilutions of the wild type and the *Δaim11* mutant on fermentative (glucose) and respiratory (ethanol/glycerol or lactate) media. While fermentative growth was unaffected, the growth of *Δaim11* mutant decreased in ethanol/glycerol and lactate compared to the WT strain (**Fig 1A**). The defect was more pronounced on lactate than on ethanol/glycerol. Unlike ethanol or glycerol, lactate donates its electrons directly to cytochrome *c* (Lodi & Ferrero, 1993), which in turn donates them to CIV, suggesting that a target of Aim11 was CIV.

**Figure 1.**
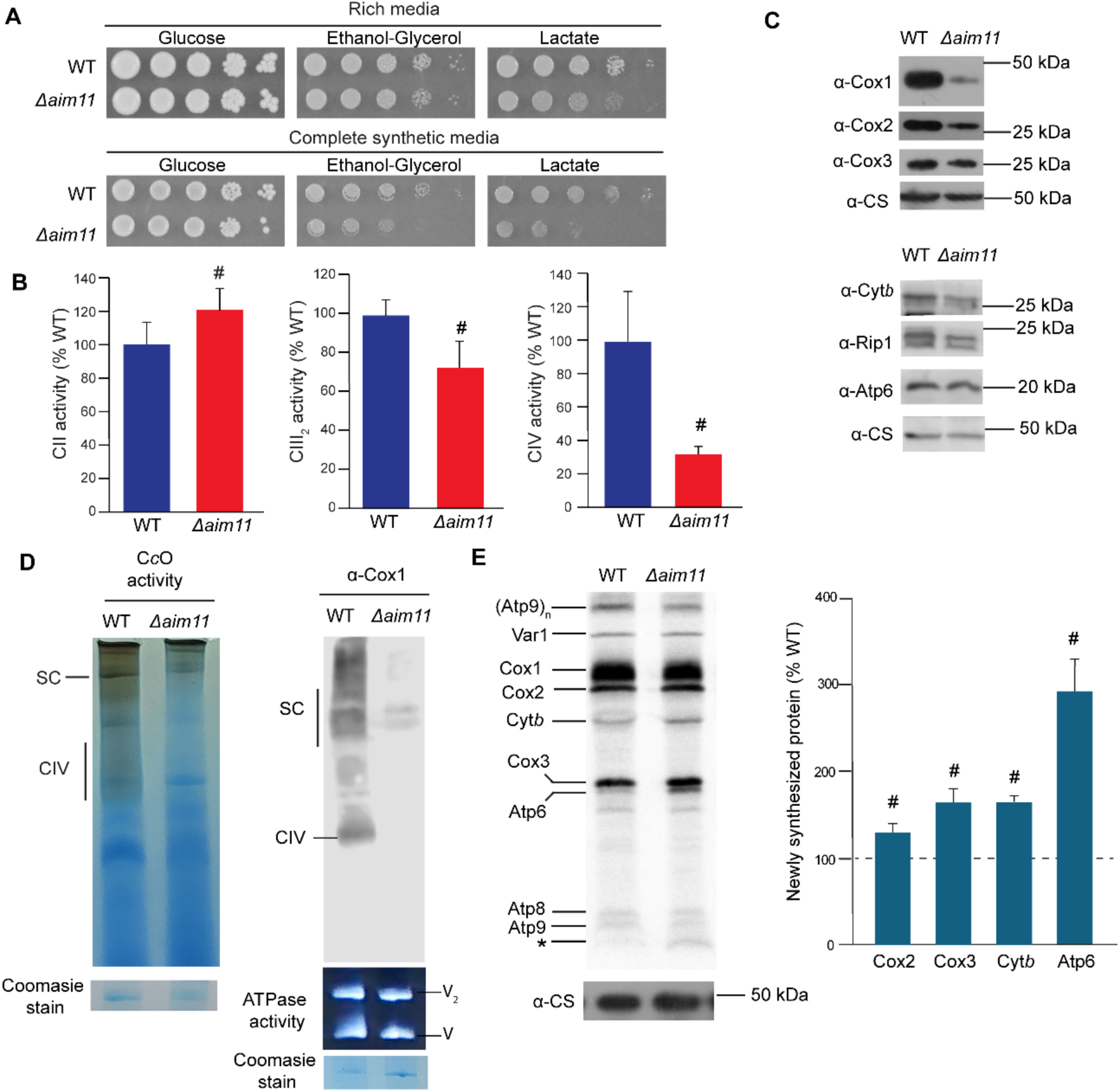
Complex IV biogenesis is decreased in the absence of Aim11. **(A)** Serial dilutions of WT and *Δaim11* strains in fermentative (glucose) and respiratory (ethanol-glycerol or lactate) media were grown for 3 days (rich media) or 5 days (complete synthetic media) at 30 °C. **(B)** Enzyme activities of CII, CIII_2_, and CIV in isolated mitochondrial membranes. Specific inhibitor-sensitive activities are represented as a percentage of the WT activity and were corrected to their respective protein content. **(C)** Western blot of mitochondria (50 µg) from the indicated strains. Citrate synthase was used as a loading control. **(D)** Mitochondria (100 µg) were solubilized with digitonin and separated by BN-PAGE. Part of the gel was transferred to a PVDF membrane for Western blot analysis, and the rest was used for in-gel staining activity of CIV and ATP synthase. **(E)** Whole-cell [^35^S]-methionine labeling of mitochondrial encoded proteins. Cox2, Cox3, Cyt*b*, and Atp6 signals were normalized to Cox1 and expressed as the percentage of wild-type. # Significant difference (*p*<0.05). * Unknown translation product present in the *Δaim11* mutant.

Next, we evaluated the activity of the respiratory complexes in isolated mitochondria. The most affected complex was CIV, where its activity was reduced to ∼30% of the WT activity (**Fig 1B**). Complex III_2_ (CIII_2_) activity was ∼73% of the WT strain, while complex II (CII) activity was slightly higher in the *Δaim11* mutant (∼120% of the WT strain). Therefore, the respiratory growth defect in the *Δaim11* mutant was due to a major reduction in CIV activity and, to some extent, also to a reduced CIII_2_ activity. We asked whether the absence of Aim11 affected CIV function or assembly. Thus, we analyzed the steady state levels of CIV subunits (Cox1-3), CIII_2_ (Cyt*b*, and Rip1), and ATP synthase (Atp6) in WT and *Δaim11* mitochondria (**Fig 1C**). Cox1 levels decreased in the mutant, as well as Cox2. Cyt*b* and Rip1. Cox3 and Atp6 subunit levels showed no significant difference between the WT and *Δaim11* strains. So far, the primary defect resulting from the absence of Aim11 has been observed in CIV activity and in the levels of Cox1 and Cox2. We wondered if the absence of Aim11 could modify the distribution of CIV between the monomeric enzyme and supercomplexes. We performed blue native electrophoresis (BN-PAGE) of digitonin-solubilized mitochondria and analyzed the CIV content by in-gel activity staining and western blot. Both immunodetection of Cox1 and in-gel CIV activity staining revealed lower levels of monomeric CIV and respiratory super complexes in the *Δaim11* strain (**Fig 1D**). In contrast, no difference was found on ATP synthase activity staining **(Fig 1D lower panel)**, suggesting that ATP synthase does not seem to contribute to the lower respiratory growth observed in **Fig 1A**. Our results indicate that the reduced activity of CIV in the *Δaim11* mutant was due to lower assembly rather than to a regulatory effect on CIV activity.

Lower levels of mitochondrially-encoded CIV subunits in the *Δaim11* mutant (**Fig 1C**) could reflect a reduced stability or diminished synthesis of the proteins. To answer this, we tested mitochondrial *de novo* protein synthesis by incubating yeast cells with [^35^S]-methionine and cycloheximide to inhibit cytosolic translation. Mitochondria were isolated, separated by SDS-PAGE, and analyzed by autoradiography. While Cox1 and Var1 synthesis were unaffected, synthesis of Cyt*b*, Cox2, and Cox3 were slightly increased (**Fig 1E**). Surprisingly, *de novo* synthesis of Atp6 greatly increased when Aim11 was absent (**Fig 1E**). However, neither Atp6 steady state levels (**Fig 1C**) nor ATP synthase levels (**Fig 1D**) reflected this increase in translation. Additionally, we consistently observed the presence of an unknown low molecular mass polypeptide that was detected in *Δaim11* cells but not in WT cells (**Fig 1E**). Taken together, these results indicate that Aim11 is required for optimal CIV assembly, while translation of mtDNA-encoded proteins was not diminished. Moreover, stability of Cox1 was found to be compromised after *AIM11* deletion.

### Aim11 affects the three assembly modules of CIV

To gain insights into the possible role of Aim11 on CIV assembly, we carried out a complexome analysis of WT and *Δaim11* mitochondria. This mass spectrometry-based technique helps identify the components of both mature mitochondrial protein complexes and their stable sub-assemblies(Cabrera-Orefice *et al*, 2022; Heide *et al*, 2012). Digitonin-solubilized mitochondria were separated by BN-PAGE. Then, each gel lane was cut into 48-60 slices and subjected individually to trypsin digestion. Identification of the peptides present in each fraction was done by LC-MS/MS. The resulting data provided a global understanding of the composition and migration of the mitochondrial proteins in the native gel and revealed the effects of the *Δaim11* mutation.

In the WT strain, Aim11 comigrated with various-sized complexes (ranging from ∼50 to 400 kDa). Its highest abundance was found at ∼150 kDa **(Fig 2A).** To establish a direct role of Aim11 in CIV assembly, we analyzed the abundance and migration patterns of the OXPHOS complexes in the WT and the *Δaim11* strains (**Fig 2B, Fig EV1A**). The migration patterns of most CII, CIII_2_, and ATP synthase subunits showed no relevant modification upon *AIM11* deletion **(Fig EV1A)**. In the mutant, the relative abundance of CIII_2_ subunits was slightly reduced, while the relative abundance of CII subunits showed a minor increase. In contrast, several CIV subunits exhibited changes in their migration patterns and relative abundance in the absence of Aim11**(Fig 2B)**. Overall, the average signal of CIV subunits in supercomplexes (∼719-950 kDa) decreased ∼70% in the mutant, whereas no major change in the monomeric fraction (∼250 kDa) was observed (as determined from calculation of the area under the curve for each region of interest). Additionally, there was an increase in non-assembled subunits **(Fig 2B).**

**Figure 2.**
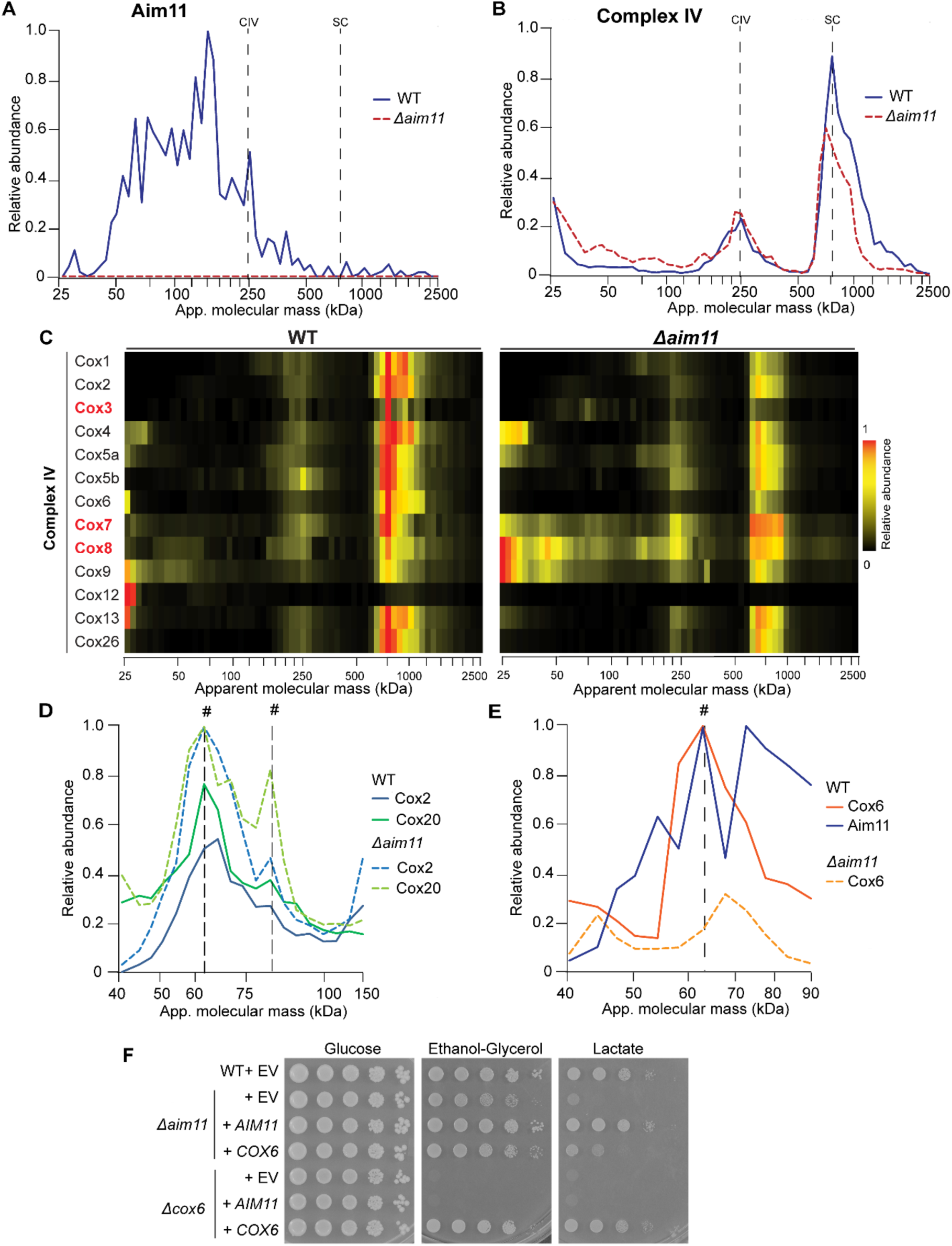
Aim11 has a role in Cox2 and Cox6 assembly. **(A)** Complexome profiling of Aim11 in mitochondria. **(B)** CIV complexome profiling in WT and *Δaim11* mitochondria. The graphic shows the average of the relative abundance of each subunit conforming CIV. **(C)** Heat maps showing the migration patterns of the individual CIV subunits in WT and *Δaim11* mitochondria. Subunits with an altered migration pattern are indicated in red characters. **(D)** Complexome profiles from Cox2 and Cox20 subassemblies showing the relative abundance from WT and *Δaim11* mitochondria. The discontinued vertical lines indicate where the proteins comigrate: # = ∼68 and ∼ 90 kDa intermediates. **(E)** Relative abundance profiles of Cox6 and Aim11 from WT and *Δaim11* mitochondria. The dotted lines indicate where the two proteins comigrate: #= ∼60 kDa. The corresponding graphs showing the averaged iBAQ values (n=2) instead of the relative abundances are shown in **Fig EV2**. **(F)** Serial dilutions of the indicated strains transformed with either empty vector (EV) or the indicated plasmids overexpressing *AIM11* or *COX6.* Cells were grown in minimum fermentative (glucose) and respiratory media (ethanol-glycerol or lactate). Growth was shown after 5 days at 30 °C.

In the WT strain, most of the CIV subunits comigrated at an apparent molecular mass of ∼250 kDa (monomeric CIV) and at ∼710 to 950 kDa (supercomplexes) **(Fig 2C)**. Cox12 was mostly found unbound in both strains, as the association with CIV seems to be destabilized under BN-PAGE conditions (Strecker *et al*, 2016). Moreover, specific subunits showed an altered migration pattern in the *Δaim11* mutant. Specifically, Cox3, Cox7 and Cox8 were accumulated in complexes with lower molecular masses (ranging from ∼50 to 150 kDa) **(Fig 2C).** These enriched low mass population indicated the accumulation of assembly intermediates.

Previous work showed that Aim11 co-purified with Cox2, Cox20 and Cox6 (Morgenstern *et al*, 2017), so we evaluated more in-depth the migration pattern of these proteins. Cox20 interacts with unassembled Cox2, and it is involved in Cox2 stabilization, translocation and processing (Elliott *et al*, 2012). We observed the Cox2-Cox20 subcomplex in two peaks (∼68 and 90 kDa) in both strains **(Fig 2D, Fig EV2A)**. In the *Δaim11* mutant, the levels of both subassemblies increased. The observed intermediates apparent molecular mass was higher than the theorical one (52 kDa), indicating that additional proteins were present. Since the migration of the Cox2-Cox20 peaks did not shift in the absence of Aim11, our data could not confirm the interaction previously reported (Morgenstern et al., 2017). Even though overall Cox2 levels were lower **(Fig 1C)**, there was an accumulation of the Cox2-Cox20 intermediates, indicating that Aim11 affects the early biogenesis steps of the Cox2 module.

Cox6, a nuclear encoded subunit that is part of the Cox1 assembly module(McStay *et al*, 2013b), was also previously reported to interact with Aim11(Morgenstern *et al*, 2017). We evaluated whether a fraction of Aim11 could migrate with Cox6. The complexome analysis showed most of Cox6 bound to CIV and supercomplexes, but only a small portion comigrated with Aim11 at ∼60 kDa **(Fig 2E, Fig EV2B).** In the *Δaim11* strain this peak disappeared, supporting the existence of a Cox6-Aim11 complex **(Fig 2E).** This association has a theorical molecular mass of 33 kDa, indicating that additional unidentified proteins must also be part of this potential complex. Altogether, the interaction previously reported (Morgenstern *et al*, 2017) and the complexome data shown here indicate that the Cox6-Aim11 interaction could help to stabilize Cox6 and facilitate its assembly into the Cox1 module. Therefore, in the absence of Aim11, Cox6 levels are expected to be reduced and, therefore, limiting for CIV assembly. To test this hypothesis, we used a 2 μ plasmid to overexpress *COX6* in different yeast strains and tested the respiratory growth of the resulting transformants. Overexpression of *COX6* resulted in a partial recovery of the respiratory growth of the *Δaim11* strain **(Fig 2F**). This result indicated that part of Aim11 role on CIV assembly is related to the assembly of Cox6, but, since respiratory growth did not fully recover, it must have additional functions (**Fig 2F**).

Next, we looked for the presence of other Cox1 module assembly intermediates that could be affected by the *Δaim11* mutation. During early assembly, newly translated Cox1 interacts with proteins like Cox14, Coa1, Coa3 and Mss51(McStay *et al*, 2013b). Additionally, the subunits Cox4, Cox5a, Cox6 and Cox8 bind to Cox1 during assembly(McStay *et al*, 2013a). Complexome profiling data showed two Cox1-containing subassemblies **(Fig 3A-B).** The first one had a molecular mass ranging from ∼90 to 100 kDa and it consisted of subunits Cox1, Cox5a and Cox8 plus the chaperones Coa1 and Cox14 **(Fig 3A, Fig EV2C)**. The calculated molecular mass of this complex is 113 kDa, which correlates well with the observed migration. The peak was sharper in the *Δaim11* strain, where Cox1, Cox5a, Cox8 and Cox14 levels increased, suggesting an accumulation of this Cox1 intermediate. The second Cox1 subassembly (∼160 kDa) contained the same components as the first subcomplex (Cox1, Cox5a, Cox8, Cox14, Coa1) plus Coa3 and Mss51 **(Fig 3B, Fig EV2D).** In the *Δaim11* mutant, there was a considerable accumulation of this subcomplex. The theoretical molecular mass of this subcomplex is 165 kDa, which also correlates well with the observed molecular mass. The accumulation of the ∼90-100 kDa and the ∼160 kDa subassemblies in the *Δaim11* mutant suggests that Aim11 assists the progression of the Cox1 module assembly.

**Figure 3.**
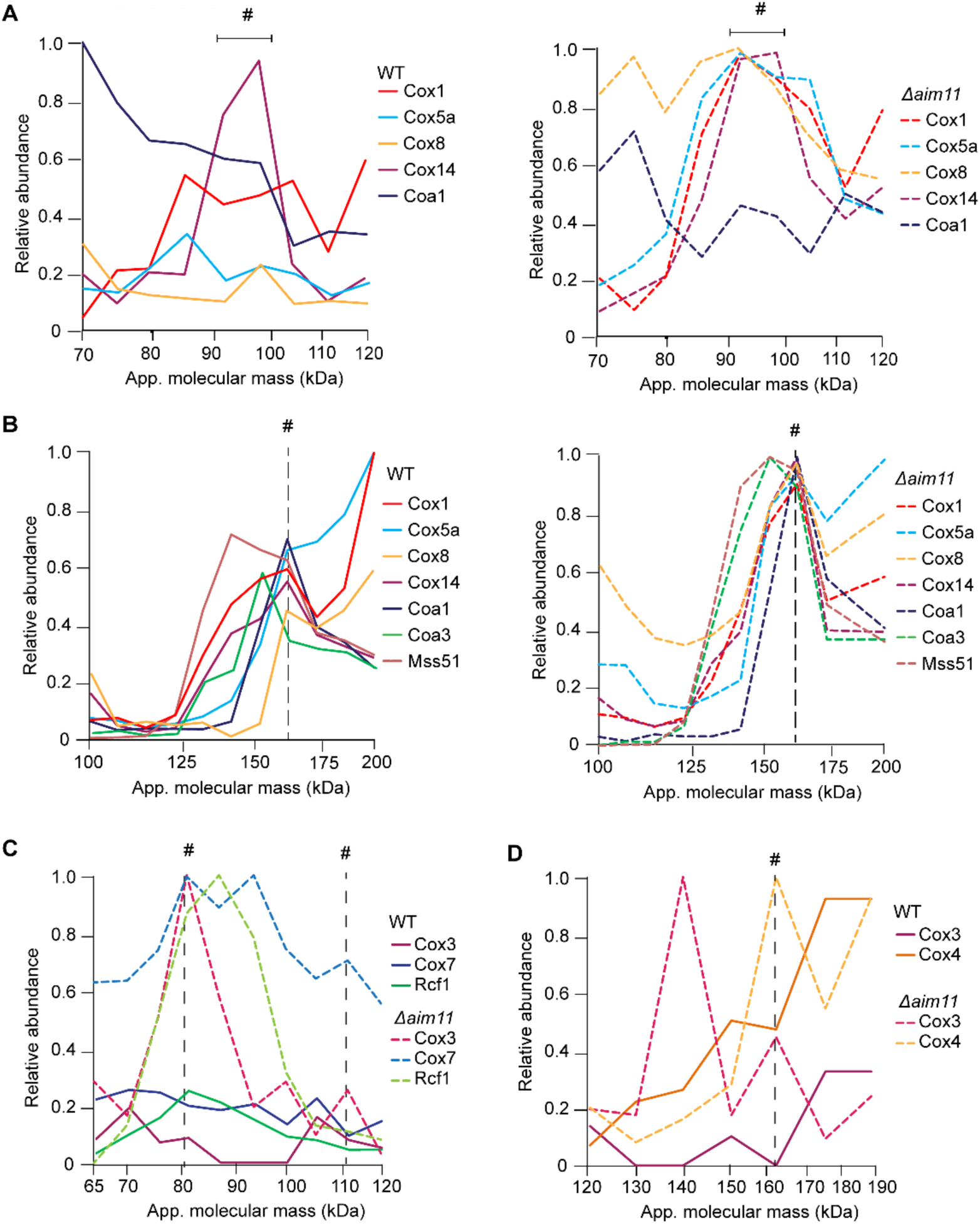
Aim11 also plays a role in the assembly of Cox1 and Cox3 modules. **(A)** Complexome analysis of Cox1 subcomplexes at ∼90-100 kDa (left panel) and *Δaim11* (right panel) mitochondria. Molecular mass interval where the proteins comigrate (#). For the corresponding graph showing average iBAQ values instead of relative values is present in **Fig EV2C**. (**B**) Relative abundance graph of the ∼160 kDa Cox1 subassembly from WT (left panel) and *Δaim11*(right panel) mitochondria. Dotted line where the proteins comigrate (#). For the corresponding iBAQ graph see **Fig EV2D**. **(C).** Complexome migration profile of Cox3, Cox7, and Rcf1 from WT and *Δaim11* mitochondria. The dotted lines indicate where the three proteins comigrated: # = ∼80 or ∼110 kDa intermediate. For the corresponding iBAQ graph see **Fig EV2E**. **(D)** Migration profile of Cox3 with Cox4 from WT and *Δaim11* mitochondria. The dotted lines indicate where the proteins comigrate: # = ∼160 kDa intermediate. For the corresponding iBAQ graph see **Fig EV2F**.

As previously stated, Cox3 also showed several new or enriched peaks in the *Δaim11* mutant (**Fig 2C**) suggesting that the Cox3 assembly module is also affected. Subunits Cox3, Cox4, Cox7, Cox13, and the assembly factor Rcf1, form the Cox3 module (Su *et al*, 2014a). While analyzing their migration patterns in the complexome dataset (**Fig 3C-D**), we detected a subcomplex with an apparent molecular mass of ∼80 kDa containing Cox3, Cox7 and Rcf1, however it was not clearly defined (**Fig 3C**). In contrast, this subcomplex was greatly enriched and defined in the *Δaim11* mutant. The theorical molecular mass of the Cox3-Cox7-Rcf1 complex was 55 kDa, indicating the association of additional unidentified proteins. In addition to the ∼80 kDa subcomplex containing Rcf1, this protein comigrated with the monomeric CIV (∼254 kDa) and with supercomplexes (∼765 kDa) (**Fig EV1B).** Meanwhile, in the *Δaim11* strain, the Rcf1 peak corresponding to the supercomplexes dramatically decreased, while the Rcf1 peak comigrating with the monomeric CIV showed a milder decrease (**Fig EV1B).** Rcf1 has a homolog, Rcf2, which is believed to be involved in super complex formation but not in Cox3 assembly (Strogolova *et al*, 2012). Unlike Rcf1, the Rcf2 migration pattern in the WT and *Δaim11* strains did not change or associated into the ∼80 kDa subcomplex (**Fig EV1B**), suggesting that Rcf2 role is independent of Aim11.

We observed two additional intermediates containing Cox3 that accumulated after *AIM11* deletion. One of them, containing Cox3 and Cox7, was detected at ∼110 kDa **(Fig 3C).** In the second one was detected at ∼160 kDa and consisted of Cox3 associated with Cox4 (**Fig 3D, Fig EV2F**). The theoretical molecular masses of both subcomplexes are 37 and 47 kDa, respectively, indicating that there must be other unidentified components in both subassemblies. These results strongly suggest that Aim11 is required for efficient progression of the Cox3 module.

Altogether, the complexome profiling analysis of WT and *Δaim11* mitochondria, indicated that Aim11 has a role in CIV biogenesis under the tested conditions. In its absence, the assembly of CIV is inefficient, resulting in an accumulation of intermediates from all three assembly modules.

### Aim11 function is facilitated by Iai11, Gep7 and Mtc3

Next, we asked whether Aim11 acts alone in CIV biogenesis or as part of a complex. Previous work has shown that Aim11 co-purifies with other uncharacterized proteins, including the Aim11 paralogue, Iai11, plus the proteins Mtc3 and Gep7 (Morgenstern *et al*, 2017). Like Aim11, Iai11 is an inner membrane protein with two transmembrane helices, with both N- and C-termini exposed to the intermembrane space (Morgenstern *et al*, 2017). Both Aim11 and Iai11 co-purified with the same set of CIV and ATP synthase subunits in high-throughput experiments (Morgenstern *et al*, 2017). Mtc3 and Gep7 are membrane integral mitochondrial proteins (Huh *et al*, 2003) with a predicted mitochondrial targeting sequence and one transmembrane helix. Like *AIM11*, the *GEP7* gene shows a negative genetic interaction with genes encoding prohibitins (Osman *et al*, 2009). The *Δmtc3* mutant is synthetically sick when it is in combination with the *cdc13-2* allele, encoding a mutant version of Cdc13, a telomeric single-stranded DNA binding protein (Addinall *et al*, 2008). Other than this, the functions of Iai11, Gep7, and Mtc3 remain unknown.

We asked if Iai11, Gep7, and Mtc3 might also be involved in CIV biogenesis. Thus, we analyzed the *Δiai11*, *Δgep7,* and *Δmtc3* single mutants, as well as double mutants to evaluate possible genetic interactions between them and *AIM11*. The *Δiai11*, *Δgep7,* and *Δmtc3* single mutants showed decreased respiratory growth as compared to the WT cells (**Fig 4A**). However, these mutants grew slightly better compared to the *Δaim11* single mutant. The double mutants showed the same phenotype as the *Δaim11* single mutant. These results suggest that Iai11, Gep7, and Mtc3 are necessary but not essential for optimal respiratory growth. Additionally, the absence of Aim11 leads to a more severe respiratory defect than the individual deletion of *IAI11*, *GEP7,* or *MTC3*.

**Figure 4.**
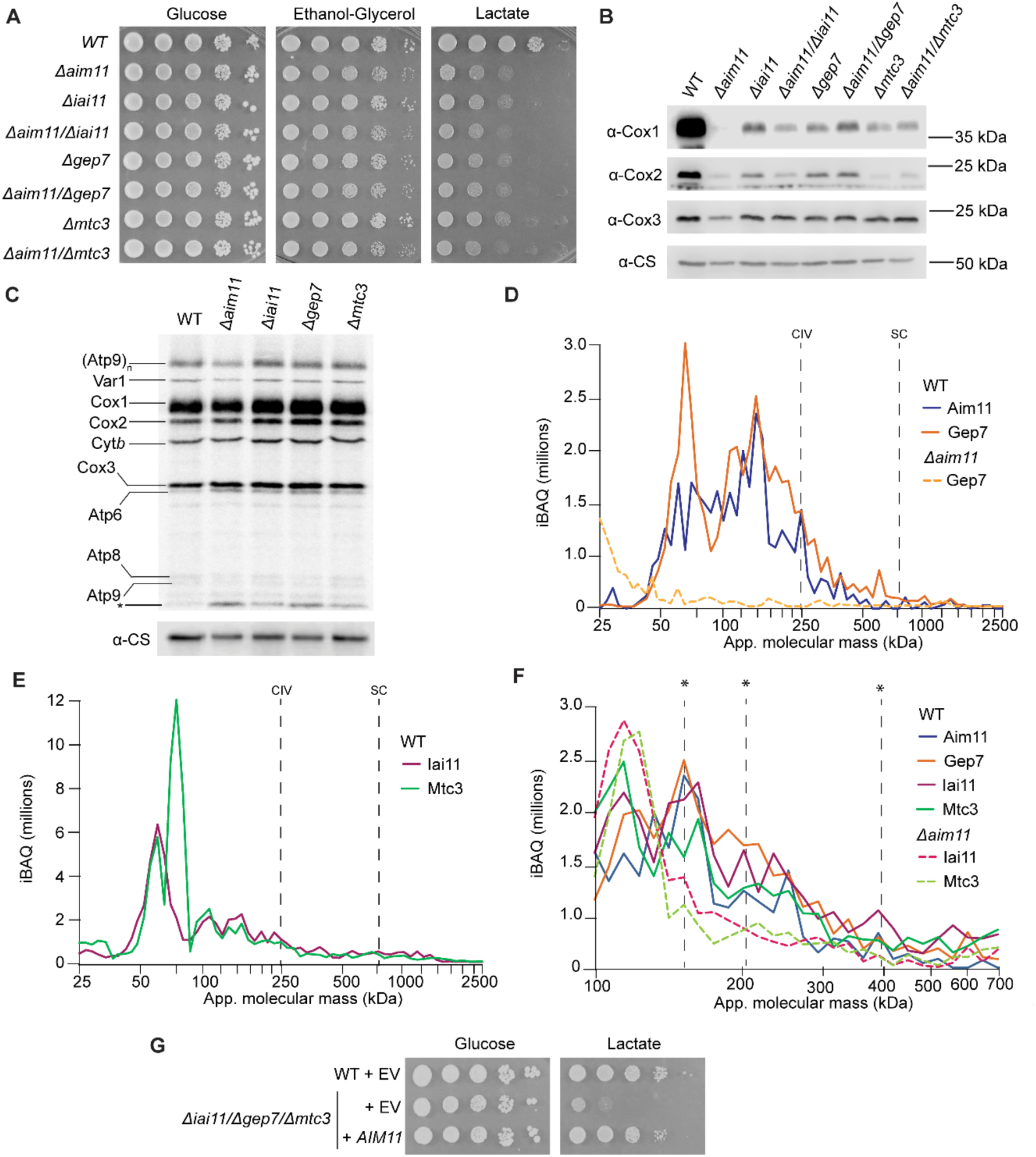
Aim11 forms the AMIGa complex, together with Iai11, Gep7, and Mtc3 that participate in cytochrome *c* oxidase biogenesis. **(A**) Serial dilutions of the indicated strains in fermentative (glucose) and respiratory media (ethanol-glycerol or lactate). Growth in CSM media was shown after 3 days. **(B)** Western blot of mitochondria (50 µg of protein per lane) showing steady state levels of the indicated proteins. CS= loading control. **(C)** Mitochondrial translation products from whole cells [^35^S]-methionine labeling on the indicated strains. An unknown translation product was present in all the null mutants (*). **(D)** Gep7 and Aim11 migration patterns in a WT and *Δaim11* mutant. Data are shown as iBAQ values. **(E)** Mtc3 and Iai11 complexome profiling in a WT strain. Data are shown as iBAQ values **(F)** Aim11, Iai11, Mtc3, and Gep7 migration patterns in a WT strain and *Δaim11* mutant. Peaks where all four proteins comigrate are indicated with an asterisk (*). Data are shown as iBAQ values. **(G)** Serial dilutions of the WT strain and *Δiai11/Δgep7/Δmtc3* triple mutant with the indicated plasmid in fermentative (glucose) and respiratory media (lactate). EV= Empty vector.

Next, we evaluated steady state levels of mitochondrial CIV subunits. Just like the *Δaim11* mutant, the absence of Gep7, Iai11, and Mtc3 showed lower levels of CIV subunits. Once again, Cox1 was the most affected subunit, although Cox2 showed a significant decrease. The Cox3 levels were only slightly decreased **(Fig 4B).** As observed for the *Δaim11* strain, in *Δiai11*, *Δgep7,* and *Δmtc3* mutants the [^35^S]-methionine labeling was not affected, except for Atp6 synthesis, which was increased compared to the WT strain (**Fig 4C**). The same unknown low molecular mass peptide was observed in all the mutants (see also **Fig 1E**). These results suggest that Aim11, Gep7, Mtc3 and Iai11 are on the same pathway during CIV assembly.

To gain a deeper understanding of Aim11 physical and functional relation with Iai11, Gep7, and Mtc3, we analyzed and compared the migration patterns of each protein in the complexome dataset. On one side, Gep7 showed a very similar migration pattern to Aim11(**Fig 4D**) but was mostly undetectable in the *Δaim11* strain (**Fig 4D**). This suggests that the stability of Gep7 depends on the presence of Aim11 and that both proteins interact. A portion of Gep7 comigrates with the Aim11-Cox6 subcomplex (**Fig EV3B**). The theoretical mass of the Aim11-Gep7-Cox6 complex (∼60 kDa) matched the observed molecular mass, which supports the idea that Aim11 and Gep7 form a heterodimer.

Iai11 and Mtc3 presented a similar migration pattern, suggesting that both proteins form a second heterodimer (**Fig 4E**). Whereas Gep7 levels were compromised in the *Δaim11* strain, Iai11 and Mtc3 levels were not significantly affected **(Fig EV3A).** Previously, it has been proposed that Aim11, Iai11, Gep7 and Mtc3 form a complex(Morgenstern *et al*, 2017). The sum of the molecular mass of one copy of each protein is ∼87 kDa, so we searched for a possible comigration at this range. We found three peaks (∼150, 200 and 400 kDa) where all four proteins were present **(Fig 4F).** In this portion, when Aim11 (and therefore Gep7) were absent, Iai11 and Mtc3 levels were greatly reduced **(Fig 4F)**, confirming that, as previously reported (Morgenstern *et al*, 2017), Aim11 forms a complex with Iai11, Gep7 and Mtc3. We named it AMIGa (Aim11-Mtc3-Iai11-Gep7 association) complex.

All four mutants share similar phenotypes, which indicate that all components of the complex are necessary for its function. We tested if the overexpression of one component of the putative Aim11-Iai11-Gep7-Mtc3 complex could compensate for the absence of another. Surprisingly, the overexpression of *AIM11* compensated for the respiratory growth defect of the single *Δgep7, Δiai11,* and *Δmtc3* mutants as well as the triple *Δgep7/Δiai11/Δmtc3* mutant (**Fig 4G**, **Fig EV3C**). In contrast, the absence of Aim11 was not compensated by overexpression of *GEP7*, *IAI11*, or *MTC3*. This result suggests that the leading component in the complex is Aim11, while Gep7, Iai11, and Mtc3 play an accessory role in CIV biogenesis. Moreover, the growth phenotype indicated that these proteins share functions, although they do not seem to have overlapping roles. Even though Gep7 depended on Aim11 to form the heterodimer **(Fig 4D)**, *AIM11* overexpression could compensate for the *Δgep7* strain respiratory growth, suggesting that Aim11 might act independently of the Aim11-Gep7 heterodimer. Finally, to reinforce the idea that Gep7, Iai11 and Mtc3 also have a common role in CIV assembly, we tested if overexpression of *COX6* could also compensate the respiratory capacity of mutant cells lacking each one of these proteins. As observed for the *Δaim11* mutant, overexpression of *COX6* partially recovered the respiratory phenotype of single *Δgep7, Δiai11 or Δmtc3* mutants (**Fig EV3D**), as well as of the triple *Δgep7/Δiai11/Δmtc3* and the quadruple *Δaim11/Δgep7/Δiai11/Δmtc3* mutants (**Fig EV3E**). This observation confirms the idea that at least a fraction of each of these four proteins contribute to the efficient assembly of CIV.

### TMEM242 is the Aim11 human orthologue

We asked whether the AMIGa complex is conserved. By primary structure analysis (BLAST), we found that Aim11, Iai11, Gep7, and Mtc3 have orthologues only among fungi but not among metazoans. We then performed a secondary structure analysis of these proteins using the HHPRED platform (Zimmermann *et al*, 2018) to search for possible structural orthologues in humans, *Drosophila melanogaster,* and *Caenorhabditis elegans*. Surprisingly, Gep7 and Mtc3 showed a similar secondary structure to human, fly, and worm COX4 subunit (which is equivalent to the yeast Cox5a/b orthologue) with a probability of >99%, and an E-value <2.5e^-17^. The similarity in secondary structure between Cox5a/b and Gep7, as well as Mtc3, could facilitate the interaction of Gep7 and Mtc3 with CIV subunits during assembly.

Next, we detected that the secondary structures of Aim11 and Iai11 were like those of the TMEM242 protein in humans, flies, and worms (probability >99%, E-value <1.9e^-23^). TMEM242 is an inner mitochondrial membrane protein implicated in complex I and ATP synthase assembly (Carroll *et al*, 2021). Alignment of the Aim11 and TMEM242 amino acid sequences indicated a low identity between them of 21.14% (**Fig 5A**). Secondary structure predictions of Aim11 and TMEM242 revealed that both proteins possess two transmembrane domains (**Fig 5B**). However, experimental data suggested that Aim11 and Iai11 N- and C- termini are exposed to the intermembrane space (Morgenstern *et al*, 2017), while exposed to the matrix in the case of TMEM242 (Carroll *et al*, 2021).

**Figure 5.**
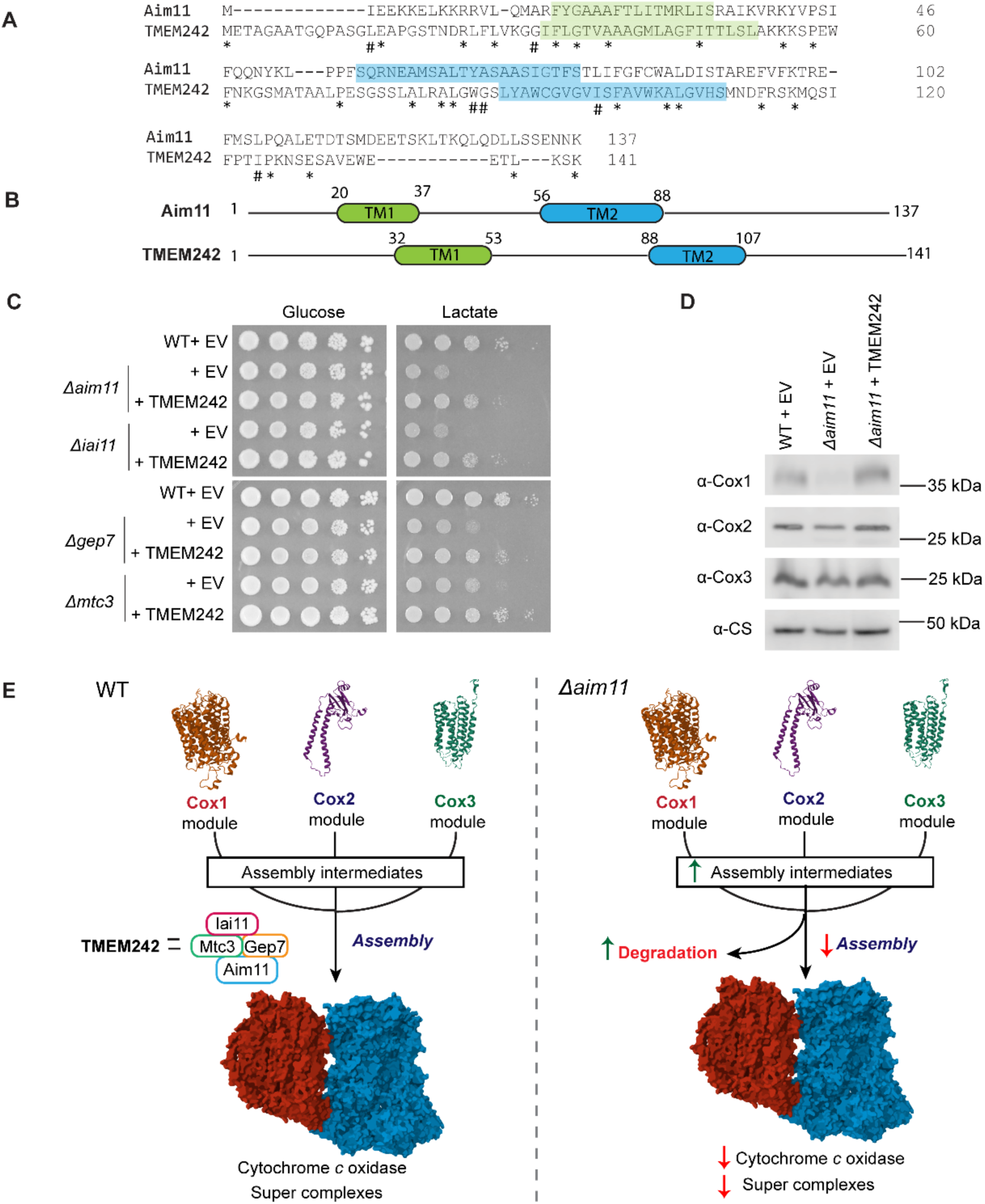
TMEM242 is the human structural orthologue of Aim11. **(A)** Alignment by CLUSTAL0 of Aim11 and its human orthologue TMEM242. Identical (*) or similar (#) amino acid residues are indicated. Predicted transmembrane domains are shown in blue or green boxes. **(B)** Secondary structure comparison between Aim11 and TMEM242. **(C)** Serial dilution of the indicated strains with overexpression of TMEM242. Growth after 5 days in respiratory (lactate) and fermentative media (glucose) at 30 °C. **(D)** Immunodetection of mitochondrial proteins from yeast cells where TMEM242 was overexpressed. CS= loading control. EV= Empty vector **(F)** Proposed model of action of the AMIGa (Aim11-Mtc3-Iai11-Gep7 association) complex on cytochrome *c* oxidase biogenesis. This complex allows the correct assembly of the Cox1, Cox2 and Cox3 modules with the rest of CIV. In its absence, intermediates from all three modules accumulated. Cox1 and Cox2 modules are degraded, resulting in lower levels of CIV and super complexes. TMEM242 is the Aim11 orthologue and can complement the absence of Aim11. 3D structures were generated based on the structure reported in (Hartley *et al*, 2020) (PDB 6T0B).

Despite the low sequence identity, we tested whether TMEM242 could compensate for the absence of different components of the Aim11-Iai11-Gep7-Mtc3 complex. We cloned the human intronless *TMEM242* from the K562 cell line into a 2 μ yeast expression vector under the *ADH* promoter and terminator. Then, the plasmid was transformed into different yeast mutants. TMEM242 compensated for the respiratory growth in the *Δaim11*, *Δgep7*, *Δiai11,* and *Δmtc3* mutants (**Fig 5C**). We then analyzed the steady state levels of CIV mitochondrial-encoded subunits in WT and *Δaim11* cells and found that TMEM242 restored Cox1 and Cox2 to WT levels (**Fig 5D**). These results strongly indicate that TMEM242 is the human orthologue of yeast Aim11.

*TMEM242* null mutant cell lines exhibited reduced levels of CIV and CIII_2_ complexes (Carroll *et al*, 2021), which is consistent with the observed phenotype of a *Δaim11* mutant, where CIV activity was dramatically reduced and complex III_2_ was slightly reduced (**Fig 1C**). Our results indicate that Aim11 modulates CIV assembly, although it is not indispensable, as 30% of CIV activity is preserved after *AIM11* deletion. This observation suggests that other proteins can compensate for the absence of Aim11. Alternatively, there could be conditions different from those used in the present study where Aim11 acquires more relevance in the assembly of CIV.

This work shows that in the absence of Aim11, assembly intermediates for all three CIV assembly modules accumulated **(Fig 5E).** Cox2 was found in complex with Cox20 (∼68 and 90 kDa) **(Fig 2D).** While Cox3 was associated with Rcf1 and Cox7 (∼80 kDa), only to Cox7 (∼110 kDa), or only to Cox4 (∼160 kDa) **(Fig 3E-F).** Cox1 was found enriched in two subassemblies (∼90-100 kDa and ∼160 kDa), both containing Cox5a, Cox8, Cox14 and Coa1. The bigger subassembly also had Coa3 and Mss51. Undoubtedly, Cox1 levels were the most affected **(Fig 1C)**, but, unlike Cox2 and Cox3, *de novo* translation of Cox1 did not increase in the null mutant, therefore, it could not offset the assembly defect **(Fig 1E).** Cox6 was not found in either subassembly, although the Aim11-Gep7 dimer could stabilize free Cox6 needed to continue Cox1 assembly **(Fig 2D, Fig EV3B)**.

In summary, Aim11 appears to exert its function on CIV assembly as part of the AMIGa complex, in which interactions among its components were previously detected (Morgenstern *et al*, 2017). While Aim11 is the main assembly factor, Iai11, Gep7, and Mtc3 function as accessory factors, since the overexpression of *AIM11* complemented the absence of the other three proteins.

## ACKNOWLEDGEMENTS

UPD is a SECIHTI fellow (CVU 883299). This project was supported by UNAM-PAPIIT (IN223623 and IN220026 for XPM, IN207023 for DGH). ACO is supported by the Netherlands Organization for Health Research and Development (ZonMW TOP-Grant 91217009) and SFB1531 (project S01#456687919) from DFG Germany. We thank Dr. Daniel Flores-Mireles for preliminary observations. We thank Dr. Ariann Mendoza-Martínez and Jana Meisterknecht for the technical help. We thank Prof. Ulrich Brandt and Dr. Ilka Wittig for providing access to mass spectrometry instrumentation. We thank Dr. Gabriel del Rio and Dr. Teresa Lara for the donation of strains. Special thanks to Dr. Mayra Furlan-Magaril, Dr. Rosario Pérez-Molina and M. C. Ayerim Esquivel-López for the donation of K562 RNA. We appreciate the donation of plasmids to Dr. Kevin Stuhl, Dr. Martin Ott and Dr. Soledad Funes. We thank Dr. Thomas D Fox, Dr. Rosemary Stuart and Dr. Soledad Funes for the gift of antisera. We thank Laura Ongay-Larios, Guadalupe Codiz-Huerta and Minerva Mora-Cabrera, the Computer Unit and Maintenance Unit of the Instituto de Fisiología Celular, UNAM. This manuscript is part of the doctoral dissertation of Ulrik Pedroza-Dávila from the Programa de Maestría y Doctorado en Ciencias Bioquímicas, Universidad Nacional Autónoma de México.

## AUTHOR CONTRIBUTIONS

U Pedroza-Dávila: conceptualization, formal analysis, investigation, validation, methodology, visualization, writing-original draft, writing–review and editing. Y. Camacho-Villasana: investigation, project administration, validation and writing-review & editing. M. Lutikurti: investigation, writing-review editing. M. Vázquez-Acevedo: investigation and writing-review & editing. D. González-Halphen: funding acquisition, resources and writing-review & editing. A. Cabrera-Orefice: data curation, formal analysis, methodology, investigation, funding acquisition, methodology, resources and writing -review editing. X Perez-Martinez: conceptualization, formal analysis, funding acquisition, methodology, project administration, resources, supervision, writing-original draft, and writing–review and editing.

## MATERIAL AND METHODS

### Yeast strains and growth conditions

Unless indicated otherwise, all *S. cerevisiae* strains are derived from the BY4742 background (ATCC 4040004). The strain list used in this study is supplementary Table 1. The yeast cells were grown in rich media (2% peptone and 1% yeast extract) or synthetic media (CSM or specific drop-out media (Formedium)) with different carbon sources (2% glucose, 2% galactose, 3% ethanol-glycerol or 2% lactate pH 5.5). For solid media 2-2.2% agar was added.

Yeast transformation was performed using the lithium acetate method. G418 resistant mutant strains are from the YKO Matα strain collection (Open Biosystems). *kanMX4* cassettes were obtained from these mutants by PCR. Otherwise, the cassettes were amplified by PCR using pUG72, pUG73, pAG25, pAG32 or pAG60 plasmids as templates (Gueldener, 2002; Goldstein & McCusker, 1999).

For C-terminal epitope tagging of the proteins of interest, a *cis*-recombination approach was used as previously reported (Schneider *et al*, 1995; Moqtaderi & Struhl, 2008). The *3xtag-URA3-3xtag* construction was amplified using the pMPY-3xHA, ZM467 or ZM474 plasmids as templates. The sequences of the oligonucleotides used are listed in supplementary Table 2.

### Plasmids construction

A list of plasmids used in this study is listed in supplementary Table 3. For yeast overexpression plasmids, the corresponding gene was amplified ∼300 bp upstream and downstream of the coding sequence by PCR. The PCR products were ligated in the pGEM-T easy vector (Promega). Plasmids were digested with *EcoRI* or *SphI/SalI* and ligated to the YEp352 2 µ plasmid (Hill *et al*, 1986).

For the *TMEM242* overexpression plasmid, cDNA was obtained from RNA of K562 human leukemia cell line (kindly donated by Dr. Mayra Furlan-Magaril lab). Then, *TMEM242* coding sequence was amplified by fusion PCR to eliminate a *HindIII* site present at 309 bp from the *AUG* start codon. A FLAG epitope was added at the C-terminal end. The construction was digested with *NotI* and *HindIII* and ligated into the pVT100U-GFP vector (Westermann & Neupert, 2000). Expression of the gene was under the *ADH* promoter and terminator. All plasmids were sequenced using specific primers.

### Mitochondrial proteins analysis

Yeast cells were obtained in logarithmic phase (O.D. 1.0 ±0.2) in galactose media and mitochondria were isolated as previously reported (Diekert *et al*, 2001). Briefly, cells were treated with zymolyase 20 T (AMSBIO 120491-1). Afterward, spheroplast were lysed by mechanical homogenization and crude mitochondria were obtained by differential centrifugation. Mitochondrial protein was quantified by the Lowry method (Markwell *et al*, 1978). Steady state levels were evaluated by SDS-PAGE (12%) and western blot using the indicated antibodies. Digitonin (Sigma D141)-solubilized mitochondria were used to separate mitochondrial complexes by BN-PAGE (4-12% gradient) (Wittig *et al*, 2006). The gel was used for cytochrome *c* oxidase or ATP synthase activities (see below) or transferred to a PVDF membrane for immunodetection. Primary antibodies used in this study are in supplementary Table 4.

### In**-**gel cytochrome *c* oxidase and ATP synthase activity

For cytochrome *c* oxidase activity staining: the gel was placed in PBS buffer pH 7.4 and horse cytochrome *c* (10 µg) and diaminobenzidine (50 µg) were added. The reaction was light protected and incubated for 2 h in agitation. The reaction was stopped and distained by addition of 10% methanol, 50% acetic acid (Wittig *et al*, 2007). For ATPase activity, the gel was placed in an incubation buffer (270 mM glycine, 35 mM Tris pH 8.4) for 3 h. The buffer was replaced with activity buffer (270 mM glycine, 35 mM Tris pH 8.4, 8 mM ATP, 14 mM MgSO_4_ ,0.2% Pb(NO_3_)_2_) and incubated for several hours until a white precipitate was formed (Wittig *et al*, 2007).

### Mitochondrial translation assay

Mitochondrial-encoded protein synthesis in whole cells was carried out as previously reported (Perez-Martinez *et al*, 2003). For [^35^S]-methionine (Perkin Elmer NEG709A001MC) *in vivo* labeling, cells were grown in YPGal two overnights at 30°C. The cultures were diluted (1 mL in 10 mL of fresh media) and incubated for 3 h at 30 °C. Cells were centrifuged and washed with sterile water. The cell pellet was resuspended in a -Met drop-out media (2% galactose) and incubated for 30 min at 30 °C. Then 5 µL of cycloheximide (10 mg/mL) were added and incubated for 5 min. Labeling was initiated by adding 7 µCi of [^35^S]-methionine. After 15 min of labeling, cells were incubated in ice-water for 5 minutes and were mechanically lysed using glass beads. Mitochondrial fraction was obtained by differential centrifugation and resuspended in loading dye. The proteins were separated by SDS-PAGE and transferred into a PVDF membrane. The membrane was placed on a phosphor screen and scanned by Typhoon 8600 (GE Healthcare). ImageJ was used for densitometry analysis. *De novo* mitochondrial protein synthesis data was normalized using the Cox1 signal, and was presented as % of WT.

### Enzymatic activity assays

Respiratory complexes activity was performed as previously reported (Flores-Mireles *et al*, 2023). In brief, complex II activity was evaluated measuring DCPIP reduction, while complex III_2_ and IV activity were estimated by measuring the reduction or oxidation of cytochrome *c*, respectively. All determinations were made at 25 °C, using a protein concentration of 20 μg/mL. Specific inhibitor-sensitive activities were obtained by subtracting the inhibitor resistant activity (malonate, antimycin A or cyanide) and normalized with the protein content. The WT strain activity was considered as 100% activity. The data was analyzed by Student’s *t*-test.

### Complexome profiling

Isolated mitochondria (100 µg) were solubilized with digitonin (3 mg/ mg protein) and separated by BN-PAGE (4-16% gradient) (Wittig *et al*, 2006). Afterwards, the gel was fixed, stained, destained, rehydrated and processed for complexome profiling analysis as formerly described (Flores-Mireles *et al*, 2023). Briefly, the gel was scanned and cut into 60 or 48 identical slices. Each slice was further diced, reduced, alkylated and trypsin digested. The resulting peptides were separated by liquid chromatography (UHPLC) and analyzed by tandem mass spectrometry (MS/MS). Then the data obtained were matched in MaxQuant (v2.6.1.0) against the reference proteome of *S. cerevisiae* (UP000002311, downloaded on 08.02.2024). The relative abundance of each protein was set based on the absolute intensity quantification values (iBAQ). To compare different samples processed in independent gel runs, we aligned the complexome profiles using COPAL (Van Strien *et al*, 2019). Finally, the hierarchically clustered complexome profiles of the mitochondrial proteins of interest were analyzed and visualized as heat maps or line charts in Microsoft Excel.

### Bioinformatic analysis

For the search of Aim11 metazoan orthologues, HHPRED platform was used (Zimmermann *et al*, 2018)*. S. cerevisiae* Aim11, Iai11, Gep7 and Mtc3 protein sequences were used to mine the *Homo sapiens, Caenorhabditis elegans and Drosophila melanogaster* proteomes, using standard parameters. Human TMEM242 was searched against the yeast proteome as an inverse query. Phobius (Kall *et al*, 2007) was used to predict Aim11 and TMEM242 transmembrane domains. The sequences of yeast Aim11 (UniProt code P87275) and human TMEM242 (UniProt code Q9NWH2) were aligned using the T-COFFEE algorithm in Jalview (Waterhouse *et al*, 2009).

## DATA AVAILABILITY

The complexome data underlying Figures 2, 3 and 4 and Supplemental Figures 1,2 and 3 will be openly available through the ComplexomE profiling DAta Resource (CEDAR) (van Strien *et al*, 2021) upon peer review of this manuscript.

