## Supplemental File for "Aim11 is a novel protein involved in the assembly of mitochondrial cytochrome *c* oxidase"

**Expanded view**

Ulrik Pedroza-Dávila[1], Yolanda Camacho-Villasana[1], Miriam Vázquez-Acevedo[1], Madhurya Lutikurti[2], Diego González-Halphen[1], Alfredo Cabrera-Orefice[2,3,4] & Xochitl Perez-Martinez[1]*

[1]Departamento de Genética Molecular, Instituto de Fisiología Celular, Universidad Nacional Autónoma de México, México City, México

[2]Research Institute for Medical Innovation, Radboud University Medical Center, Nijmegen, the Netherlands

[3]Functional Proteomics Center, Goethe University, Frankfurt am Main, Germany

[4]Institute of Biochemistry, Faculty of Medicine, Justus Liebig University, Giessen, Germany

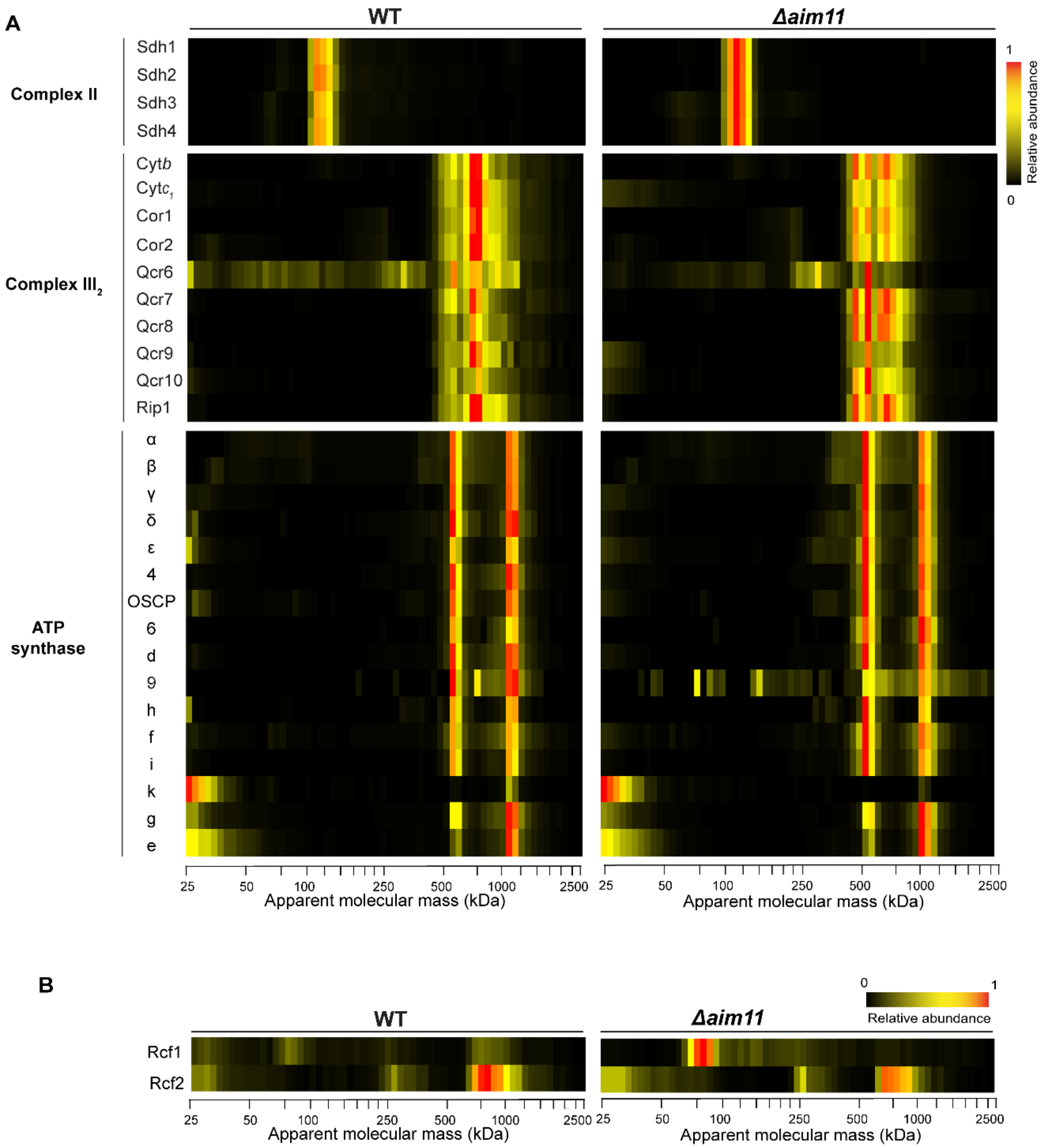


**Expanded View Figure 1. Absence of Aim11 does not alter CII, CIII$_2$ or ATP synthase levels. (A)** Heat maps of the complexome profiling analysis showing the migration patterns of CII, CIII$_2$ and ATP synthase individual subunits. **(B)** Heat maps of the complexome profiling analysis showing the migration patterns of Rcf1 and Rcf2.

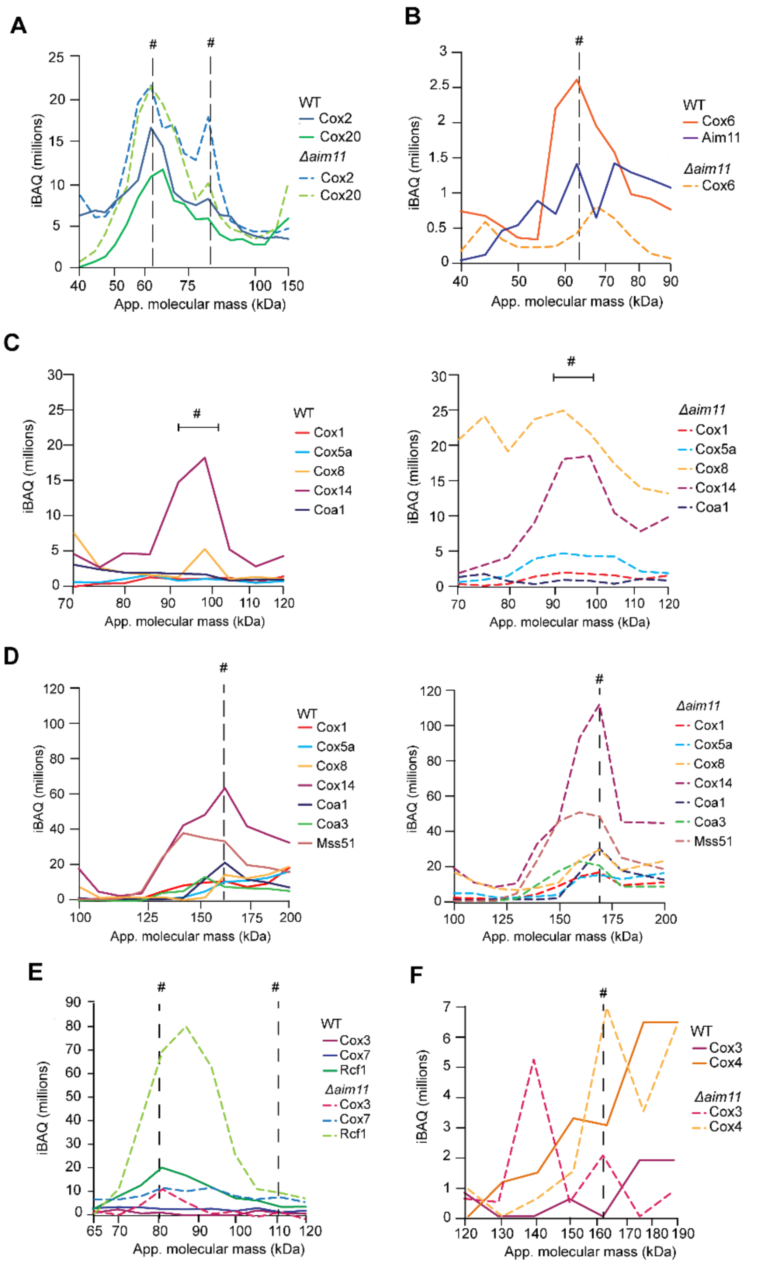


**Expanded View Figure 2. Absence of Aim11 alters the three CIV assembly modules.** Abundance and migration patterns of different CIV subunits and assembly factors. The y-axis shows averaged iBAQ values (n=2) from each protein.

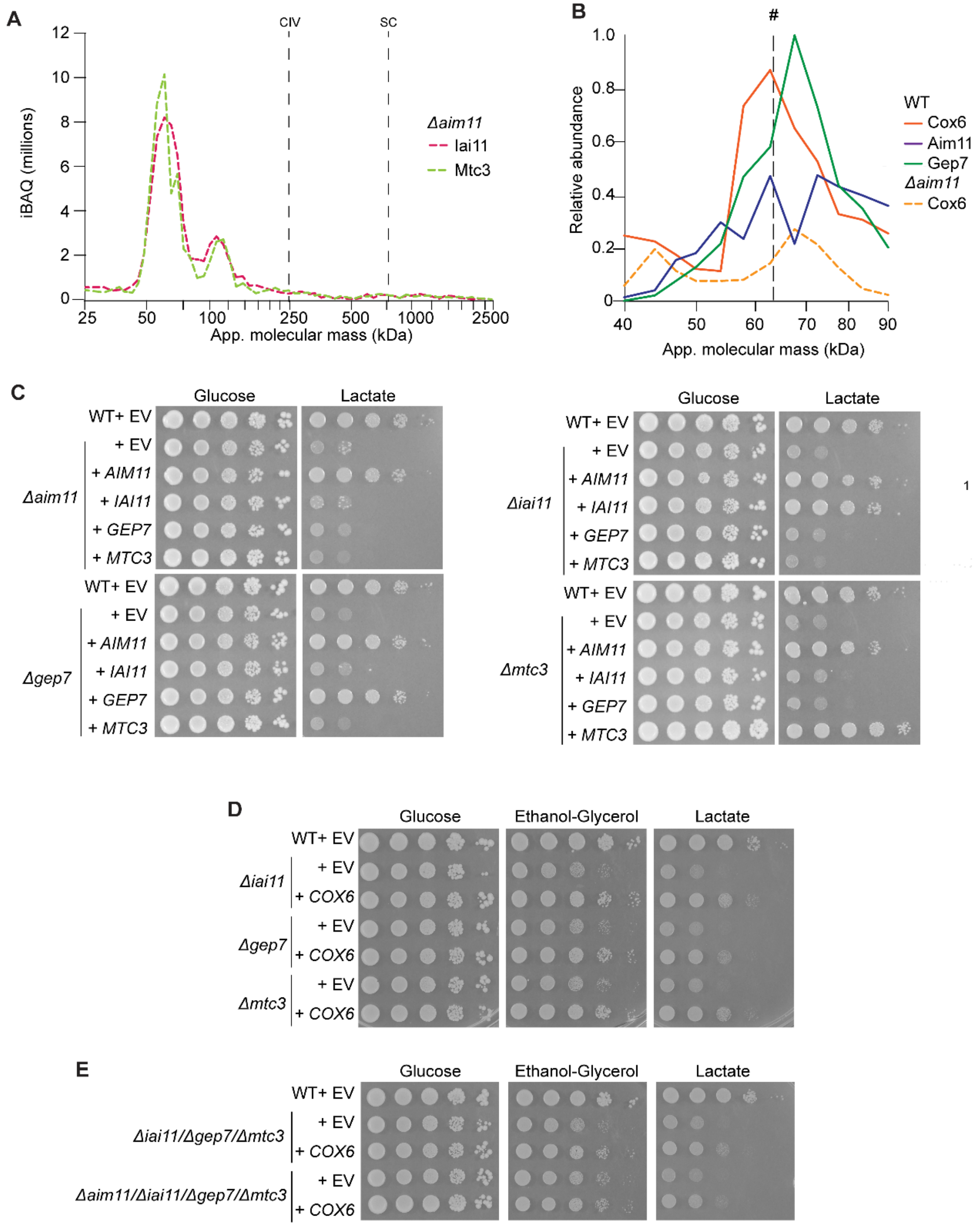


**Expanded View Figure 3. Aim11 is the core component of a complex containing Gep7, Iai11 and Mtc3 (AIMGa complex). (A)** Complexome profiles of Gep7 and Mtc3 when Aim11 is absent**. (B)** Comigration of Cox6, Aim11 and Gep7

at around 60 kDa **(C-E)** Serial dilutions of the null mutant strains with the indicated overexpression plasmids in fermentative (glucose) and respiratory media (ethanol-glycerol or lactate). Cells were grown for 5 days at 30 °C. EV is empty vector.

**Supplementary Table 1.**

| Strain | Genotype | REF |
|---|---|---|
| BY4742 | *Matα, his3-delta1, leu2-delta0, lys2-delta0, ura3-delta0, BY4742* | YKO Matα strain collection (Open Biosystems) |
| UP87 | *Matα, his3-delta1, leu2-delta0, lys2-delta0, ura3-delta0, Δaim11::LEU2, BY4742* | This study |
| UP91 | *Matα, his3-delta1, leu2-delta0, lys2-delta0, ura3-delta0, Δaim11::URA3, BY4742* | This study |
| *Δaim11* | *Matα, his3-delta1, leu2-delta0, lys2-delta0, ura3-delta0, Δaim11::kanMX4, BY4742* | YKO Matα strain collection (Open Biosystems) |
| *Δiai11* | *Matα, his3-delta1, leu2-delta0, lys2-delta0, ura3-delta0, Δiai11::kanMX4, BY4742* | YKO Matα strain collection (Open Biosystems) |
| *Δgep7* | *Matα, his3-delta1, leu2-delta0, lys2-delta0, ura3-delta0, Δgep7::kanMX4, BY4742* | YKO Matα strain collection (Open Biosystems) |
| *Δmtc3* | *Matα, his3-delta1, leu2-delta0, lys2-delta0, ura3-delta0, Δmtc3::kanMX4, BY4742* | YKO Matα strain collection (Open Biosystems) |
| *Δcox6* | *Matα, his3-delta1, leu2-delta0, lys2-delta0, ura3-delta0, Δcox6::kanMX4, BY4742* | YKO Matα strain |

| | | |
|---|---|---|
| | | collection (Open Biosystems) |
| UP85 | *Matα, his3-delta1, leu2-delta0, lys2-delta0, ura3-delta0, Δaim11::LEU2, Δiai111::kanMX4, BY4742* | This study |
| UP89 | *Matα, his3-delta1, leu2-delta0, lys2-delta0, ura3-delta0, Δaim11::LEU2, Δgep7::kanMX4, BY4742* | This study |
| UP135 | *Matα, his3-delta1, leu2-delta0, lys2-delta0, ura3-delta0, Δaim11::LEU2, Δmtc3::kanMX4, BY4742* | This study |
| UP227 | *Matα, his3-delta1, leu2-delta0, lys2-delta0, ura3-delta0, Δmtc3::kanMX4, Δgep7::natMX4, BY4742* | This study |
| UP228 | *Matα, his3-delta1, leu2-delta0, lys2-delta0, ura3-delta0, Δaim11::LEU2, Δmtc3::kanMX4, Δgep7::natMX4, BY4742* | This study |
| UP234 | *Matα, his3-delta1, leu2-delta0, lys2-delta0, ura3-delta0, Δmtc3::kanMX4, Δgep7::natMX4, Δiai11::hphMX4, BY4742* | This study |
| UP235 | *Matα, his3-delta1, leu2-delta0, lys2-delta0, ura3-delta0, Δaim11::LEU2, Δmtc3::kanMX4, Δgep7::natMX4, Δiai11::hphMX4, BY4742* | This study |
| UP120 | *Matα, his3-delta1, leu2-delta0, lys2-delta0, ura3-delta0, AIM11-3xFLAG-URA3-3xFLAG, BY4742* | This study |
| UP121 | *Matα, his3-delta1, leu2-delta0, lys2-delta0, ura3-delta0, AIM11-3xFLAG, BY4742* | This study |
| UP173 | *Matα, his3-delta1, leu2-delta0, lys2-delta0, ura3-delta0, MTC3-3xHA-URA3-3xHA, BY4742* | This study |
| UP177 | *Matα, his3-delta1, leu2-delta0, lys2-delta0, ura3-delta0, MTC3-3xHA, BY4742* | This study |
| UP178 | *Matα, his3-delta1, leu2-delta0, lys2-delta0, ura3-delta0, IAI11-3xV5-URA3-3xV5, BY4742* | This study |
| UP200 | *Matα, his3-delta1, leu2-delta0, lys2-delta0, ura3-delta0, IAI11-3xV5, BY4742* | This study |
| UP174 | *Matα, his3-delta1, leu2-delta0, lys2-delta0, ura3-delta0, GEP7-3xHA-URA3-3xHA, BY4742* | This study |

| | | |
|---|---|---|
| UP176 | *Matα, his3-delta1, leu2-delta0, lys2-delta0, ura3-delta0, GEP7-3xHA, BY4742* | This study |
| NB40 3c rho$^0$ | *Mata*, *arg8::hisG, leu2-3,112, lys2, ura3-52, his3ΔHindIII, D273-10B ($\rho^0$)* | (Bonnefoy *et al*, 2001) |

In parenthesis is the mitochondrial genotype.

**Supplementary Table 2.** Oligonucleotides used in the present work.

| Oligonucleotide | Sequence (5’-3’) | Use |
|---|---|---|
| AIM11 Fw1 | CCGAGCTCGCGGATTGATCGGTTCATGGCAATCTCATGGTCCC | Amplify *AIM11* gene ~300 bp upstream and downstream the coding sequence. |
| AIM11 Rv1 | GGCTGCAGCGCGGGCTACTCTGTTTTGCACTGTGAGGTATGCTC | |
| AIM11 Fw2 | GGATGGGGAACTTGATTCTTTGACGGGTTTGTCAGCAATTAGTACGACTATCCCCAGCTGAAGCTTCGTACGCTGC | Generate *URA3* or *LEU2* cassete for *AIM11* deletion by homologous recombination. For pUG72 or pUG73 plasmids. |
| AIM11 Rv2 | GCCGGCAACGGCTCGTAGTTATACCATTATTTACAGTTTAAAGAGATTAAGCCCCGCATAGGCCACTAGTGGATCTG | |
| AIM11 S1 | GGAATGAAGCCATGTCAGCGTTAACGTATGCATCC | Forward primer to sequence *AIM11* constructions. |
| AIM1pMPY F | ACTAACTAAGCAATTGCAAGACCTCCTGTC GAGCGAAAACAACAAGAGGGAACAAAAGCTGG | To tag Aim11 at the C-terminus. To Use pMPY-HAx3, ZM467 or ZM474 plasmid. |
| AIM11 pMPY R | TATACCATTATTTACAGTTTAAAGAGATTAAGCCAATGCGTAGTGCTACTGTAGGGCGAATTGGG | |
| TMEM242 F | GTGAAGCTTATGGAGACAGCGGGCGCTGCAACTGG | To clone TMEM242-FLAG from cDNA by fusion PCR. |
| TMEM242 P2 | CCTAAAGCCTTCCAGACTGCGAAGCTAATC | |

| | | |
|---|---|---|
| TMEM242 P3 | GATTAGCTTCGCAGTCTGGA AGGCTTTAGG | |
| TMEM242 R | ACTGCGGCCGCTCACTTGTC GTCATCGTCTTTGTAGTCCGC CGCTTTGGATTTCAATGTTTC CTCCC | |
| IAI11 F | TTGCCTCCTTGAGACACTTGT TCAAAGC | Amplify *IAI11* gene ~300 bp upstream and downstream the coding sequence. |
| IAI11 R | TCATCCGTTTGAGAAGCATAA TAGG | |
| IAI11 F2 | AGCAAGTAGTGGACAGCTTA GTTAAGACACACAATTCATCT CTTTGTAAA AGGGAACAAAAGCTGG | To tag Iai11 at the C-terminus. Use pMPY-HAx3, ZM467 or ZM474 plasmid. |
| IAI11 R2 | ATATAAGTAGTGGAATATATT ATTGAAACTACTACTATGGTA TAACACTACTGTAGGGCGAA TTGGG | |
| IAI11 S1 | GGTATATTTGGTATGGGCATC ACAGG | Forward primer to sequence *IAI11* constructions. |
| IAI11 F3 | GTAGTACTTGTCTTATCTTGC TTTTGTTCAACTGCACTTGTA AATCAGTGACCAGCTGAAGC TTCGTACGC | Generate *URA3*, *nat*MX4, *hph*MX4 or *kan*MX4 cassete for *IAI11* deletion by homologous recombination. For pAG25, pAG32, pAG60 or pAFA-*kan* plasmids. |
| IAI11 R3 | CACATATAAGTAGTGGAATAT ATTATTGAAACTACTACTATG GTATAACACGATGAATTCGAG CTCGTT | |
| GEP7 F | CCACCTCAAAGTCCAAGTAAT TGTCC | Amplify *GEP7* gene ~300 bp upstream and downstream the coding sequence. |
| GEP7 R | CATCGTGAGTTTCTTTCTCTA TCGATCC | |

| | | |
|---|---|---|
| GEP7 F2 | AACAAAAAACGAAGGAGCTGCCCGCACAGAAGAGTACTGTAATATCAGAAAGGGAACAAAAGCTGG | To tag Gep7 at the C-terminal. Use pMPY-HAx3, ZM467 or ZM474 plasmid. |
| GEP7 R2 | TGAGTGGCACTTTTAACTGCAAGAATGAAGAGTGACCCCATTTTTTTTTAACTGTAGGGCGAATTGGG | |
| GEP7 F3 | TATATACACATGTTCTCGCTAAAATCAGTTAAAAGGTCCGATTGAGCCAGCTGAAGCTTCGTACGC | Generate *URA3*, *nat*MX4, *hph*MX4 or *kan*MX4 cassete for *GEP7* deletion by homologous recombination. For pAG25, pAG32, pAG60 or pAFA-*kan* plasmids. |
| GEP7 R3 | CACATATAAGTAGTGGAATATATTATTGAAACTACTACTATGGTATAACACGATGAATTCGAGCTCGTT | |
| GEP7 S1 | CTCAATTTGTCCTGACAAACTCTTCTCC | Forward primer to sequence *GEP7* constructions. |
| MTC3 F | GGATAAATGGCTGGAAGCTCTTTGAGG | Amplify *MTC3* gene ~300 bp upstream and downstream the coding sequence. |
| MTC3 R | CTAAGAGTGACTAATGTAGGTAAGTGC | |
| MTC3 F2 | ATAATTGGAAACAAGATAAAAAGCTAGAGGAACAATTAAGGGATCTTGTAAGGGAACAAAAGCTGG | To tag Mtc3 at the C-terminus. Use pMPY-HAx3, ZM467 or ZM474 plasmid. |
| MTC3 R2 | GTACTCTGCGCAAGCGCATATATATATATACTATTGTTGCTTCCAATTTACTGTAGGGCGAATTGGG | |
| MTC3 F3 | AAGACCACCCAAAAGATATCATTTCAACATAGGTAGAAATAA | Generate *URA3*, *nat*MX4, *hph*MX4 or *kan*MX4 cassete for |

| | | |
|---|---|---|
| | GGCAGAAGCCAGCTGAAGCTTCGTACGC | *MTC3* deletion by homologous recombination. For pAG25, pAG32, pAG60 or pAFA-*kan* plasmids. |
| MTC3 R3 | AACGTACTCTGCGCAAGCGCATATATATATATACTATTGTTGCTTCCAATTCGATGAATTCGAGCTCGTT | |
| MTC3 S1 | CAAGGAATACTTACGATGGAGGATGC | Forward primer to sequence *MTC3* constructions. |
| COX6 F1 | GCCAATCAGGGCCCCGCGCGTTATTTC | Amplify *COX6* gene ~300 bp upstream and downstream the coding sequence. |
| COX6 R1 | ATATTAAAGGTAATCTGTGACCAGCCC | |
| NAT Rev | GTTGTTTATGTTCGGATGTG | Reverse primer to confirm recombination of *natMX4* cassete in the mutant. |
| HYG Rev | GCAATCGCGCATATGAAATC | Reverse primer to confirm recombination of *hphMX4* cassete in the mutant. |
| KanB | CTGCAGCGAGGAGCCGTAAT | Reverse primer to confirm recombination of kanMX4 cassete in the mutant. |
| ADH1 F1 | CGTCATTGTTCTCGTTCCCTTTCTTCC | To sequence construction in pVT-100 plasmid. |
| ADH1 R1 | GAGTCACTTTAAAATTTGTATACAC | |

**Supplementary Table 3.** Plasmids used or constructed in the present study.

| Plasmid | Relevant characteristics | REF |
|---|---|---|
| pGEM-T Easy vector | PCR cloning plasmid. *AmpR* bacteria selection marker. | Promega corportation |
| YEp352 | Yeast/*E. coli* shuttle 2 µ plasmid. AmpR bacteria selection marker. *URA3* yeast selection marker. | (Hill *et al*, 1986) |
| pVT100-GFP | Yeast expression 2 µ vector. mtGFP under *ADH* promoter. AmpR bacteria selection marker. *URA3* yeast selection marker. | (Westermann & Neupert, 2000) |
| pMPY-HAx3 | Epitope tagging template for *3xHA-URA3-3xHA*. AmpR bacteria selection marker. | (Schneider *et al*, 1995) |
| ZM467 | Epitope tagging template for *3xFLAG-URA3-3xFLAG*. AmpR bacteria selection marker. | (Moqtaderi & Struhl, 2008) |
| ZM474 | Epitope tagging template for *3xV5-URA3-3xV5*. AmpR bacteria selection marker. | (Moqtaderi & Struhl, 2008) |
| pUG72 | *loxP–URA3–loxP* template plasmid. AmpR bacteria selection marker. | (Gueldener, 2002) |
| pUG73 | *loxP–LEU2–loxP* template plasmid. AmpR bacteria selection marker. | (Gueldener, 2002) |
| pAG25 | *natMX4* cassete template plasmid. AmpR bacteria selection marker. | (Goldstein & McCusker, 1999) |
| pAG32 | *hphMX4* cassete template plasmid. AmpR bacteria selection marker. | (Goldstein & McCusker, 1999) |
| pAG60 | *URA3* cassete template plasmid. AmpR bacteria selection marker. | (Goldstein & McCusker, 1999) |
| pUP10 | *TMEM242*-FLAG in pVT100U. Under ADH promoter and terminator | This study |
| pUP17 | *AIM11-3xFLAG* in pGem-T Easy | This study |
| pUP18 | *AIM11-3xFLAG* in YEp352 | This study |

| | | |
|---|---|---|
| pUP13 | *IAI11-3xV5* in pGem-T Easy | This study |
| pUP14 | *IAI11-3xV5* in YEp352 | This study |
| pUP12 | *GEP7-3xHA* in pGem-T Easy | This study |
| pUP16 | *GEP7-3xHA* in YEp352 | This study |
| pUP11 | *MTC3-3xHA* in pGem-T Easy | This study |
| pUP15 | *MTC3-3xHA* in YEp352 | This study |
| pUP28 | *COX6* in pGem-T Easy | This study |
| pUP29 | *COX6* in YEp352 | This study |

Supplementary Table 4.- **Primary antibodies**

| Antibody | Dilution | Secondary | Source |
|---|---|---|---|
| α-Cox1 | 1: 40000 | Rabbit | (García-Villegas *et al*, 2017) |
| α-Cox2 | 1:2000 | Mouse | Mitoscience |
| α-Cox3 | 1:2000 | Mouse | Mitoscience |
| α-CS | 1:1000 | Rabbit | Gift for XPM from Dr. Thomas Fox |
| α-Atp6 | 1:2000 | Rabbit | Gift for XPM from Dr. Soledad Funes |
| α-Cyt*b* | 1:1000 | Rabbit | Gift for XPM from Dr. Rosemary Stuart |
| α-Rip1 | 1:1000 | Rabbit | Gift for XPM from Dr. Rosemary Stuart |
